# Prognostic significance of Ki-67, BAX, Caspase-3, and VEGF expression in feline intestinal lymphoma

**DOI:** 10.1101/2025.06.01.657294

**Authors:** Natalia Camargo Faraldo, Andresa da Fontoura Souza Rosenfeld, Katia Graciele Brombatti, Renee Laufer Amorim, Suzana Terumi Honda Battaglia, Renata Afonso Sobral, Carlos Eduardo Fonseca Alves

## Abstract

Feline intestinal lymphoma is the most common gastrointestinal neoplasm in cats and presents variable biological behavior and treatment responses. Despite its clinical importance, reliable prognostic biomarkers for this neoplasm are still lacking. This study aimed to investigate the immunohistochemical expression of Ki-67, BAX, caspase-3, and VEGF-A in feline intestinal lymphoma and evaluate their potential associations with overall survival and response to chemotherapy. Thirty cases of feline intestinal lymphoma were retrospectively analyzed. Clinical data, histopathological classification (lymphocytic vs. lymphoblastic), and immunophenotyping (B-vs. T-cell origin) were assessed. Immunohistochemistry was performed to evaluate the expression levels of Ki-67, BAX, caspase-3, and VEGF-A, which were classified as high or low. Associations between biomarker expression, lymphoma subtype, treatment status, and overall survival were analyzed via Kaplan-Meier curves and statistical comparisons. Most cases were of T-cell origin (22/30), with 17 classified as lymphocytic and 13 as lymphoblastic lymphoma. Chemotherapy significantly increased overall survival (p = 0.0008). Low expression of BAX (p = 0.0032), high expression of Ki-67 (p = 0.0184), and low expression of caspase-3 (p = 0.0222) were associated with reduced survival. VEGF-A expression was not significantly associated with survival (p = 0.0556), although a trend toward worse prognosis in high-expression cases was observed. When biomarker expression was analyzed in conjunction with chemotherapy status, the prognostic value of each marker was enhanced, with all the markers showing significant associations with survival (p < 0.0001). BAX, caspase-3, and Ki-67 expression levels are associated with prognosis in feline intestinal lymphoma, particularly when considered alongside treatment status. These markers may assist in predicting survival outcomes and tailoring therapeutic strategies in affected cats.

## Introduction

Lymphoma is one of the most common cancers in cats and arises from the malignant transformation of lymphocytes. This neoplasm can occur in various anatomical forms, including multicentric, mediastinal, and alimentary forms, with clinical signs varying according to the affected site. Despite advancements in diagnosis and treatment, feline lymphoma remains a complex disease with diverse presentations and prognoses, and further investigations are needed to optimize management strategies [1] [2].

Alimentary lymphoma is one of the most prevalent anatomical forms of feline lymphoma, followed by the mediastinal and multicentric forms [3]. This neoplasm primarily affects the gastrointestinal tract, involving the small or large intestine, liver, and pancreas, and is characterized by the infiltration of neoplastic lymphocytes, with or without mesenteric lymph node involvement [4]. Histopathologically, it can be classified as small-cell lymphoma, also known as lymphocytic lymphoma, or large-cell lymphoma, referred to as lymphoblastic lymphoma [4]. Lymphocyte infiltration generally occurs irregularly across the intestinal villi, potentially progressing to the submucosa and resulting in transmural involvement [4]. In some cases, epitheliotropism is observed, characterized by lymphocyte clustering among epithelial cells or diffuse infiltration within the villi and crypts of the epithelium [4]. Additionally, alimentary lymphoma presents clinical manifestations such as weight loss, vomiting, and diarrhea, reflecting gastrointestinal dysfunction [4].

Prognostic factors in feline lymphomas include anatomical location, disease stage at diagnosis, the presence or absence of FeLV and FIV infection, and the initial response to treatment [5]. Prognostic markers are measurable biological indicators that provide essential information about the probable evolution of a disease [6]. The accurate detection of prognostic molecular indicators plays a crucial role in simplifying the logical selection of promising therapeutic targets during the development of new cancer therapies. Furthermore, it enables consistent and replicable risk stratification, playing an essential role in clinical trials [7]. The Ki-67 antibody, located in the nucleus of dividing cells (G1, S, G2, M), excluding the resting phase (G0), is a cell proliferation indicator [8]. Its high expression in tumors suggests its aggressiveness and invasive potential [9]. The BAX gene and its encoded protein play pivotal roles in apoptosis, inducing mitochondrial membrane permeabilization [10]. In lymphomas, BAX expression serves as an indicator of the apoptotic response, influencing disease progression and treatment efficacy. Studies have explored BAX expression in human and animal lymphomas, employing immunohistochemical and molecular analyses to correlate BAX with cell proliferation, invasion, and therapeutic response [11]. Caspase-3, an enzyme from the caspase family, regulates apoptosis and the cellular stress response, influencing genomic integrity, tumor suppression, and intracellular signaling. Caspase-3 activity can be used to evaluate the efficacy of apoptosis in cancer cells; a reduction in this activity contributes to lymphoma progression [12]. Vascular endothelial growth factor (VEGF) is a protein that plays a key role in angiogenesis and is central to the formation of new blood vessels. Elevated levels of VEGF are associated with increased tumor vascularization, indicating the potential for aggressive growth and spread [13].

Thus, this study aimed to evaluate the prognostic value of the markers Ki-67, BAX, Caspase-3, and VEGF-A in cats with lymphoma.

## Methodology

### Sample origin

A population sampling analysis was conducted via computational software (G-power®, Brunsbüttel, Germany) with the following criteria: type I error rate (α) = 0.05, type II error rate (β) = 0.2, and a statistical power of 80%. On the basis of these analyses, we established 30 cases and the minimum number of cases. The samples were retrospectively collected from different institutions.

### Inclusion and Exclusion Criteria

The primary inclusion criterion was cats with feline lymphoma diagnosed via biopsy before any chemotherapy or corticosteroid treatment. The availability of a paraffin block with sufficient tissue for immunohistochemistry analysis was then necessary. Only cats that underwent clinical staging (TNM) and were diagnosed with alimentary lymphoma, independent of age and sex, were included in this research. Clinical information was obtained from patient records. Patients for whom treatment and survival data could not be obtained were excluded from the study. Survival was calculated on the basis of the time from lymphoma diagnosis to patient death. We censored the survival data only for patients who were alive. Overall survival was considered the time from the confirmatory diagnosis until death. The patients who were still alive were considered the time of submission of this manuscript for publication.

### Histology and immunohistochemistry

For histological analysis, paraffin-embedded samples were sectioned into 4 μm slices, placed on polished histological slides, and subjected to hematoxylin and eosin (HE) staining. The feline intestinal lymphomas were classified as lymphocytic or lymphoblastic according to previous methods [14]. The immunostaining assessment for antibodies was determined through optical microscopy. For BAX and VEGF-A, cytoplasmic expression was assessed, and for Caspase-3 and Ki67, nuclear expression was assessed. The BAX [15], Caspase-3 [16], Ki67 [17] and VEGF-A [18] clones were previously validated and tested in feline tissue. The samples were sectioned from paraffin blocks at a thickness of 4 μm and mounted on positively charged slides (Starfrost, Knittel, Bielefeld, Germany). For antigen retrieval, the slides were incubated in a high-pH solution (EnVision FLEX Target Retrieval Solution High pH 50x - Dako) in a pressure cooker (Pascal, Dako, Carpinteria, CA, USA) for 45 minutes. Endogenous peroxidase activity was inhibited via the addition of 8% hydrogen peroxide in methanol for 15 minutes, followed by treatment with 8% skim milk at room temperature. The samples were then incubated overnight at 4°C with antibodies against Ki-67 (clone MIB-1, mouse monoclonal IgG1, Dako, Santa Clara, CA, USA, 1:100), Caspase-3 (clone D175, rabbit polyclonal, Cell Signaling Technology, Danvers, MA, USA, 1:1000), BAX (rabbit polyclonal, Dako, Santa Clara, CA, USA, 1:1500), and VEGF-A (clone VG-1, mouse monoclonal IgG1, Santa Cruz Biotechnology, Dallas, TX, USA, 1:100), following standardized dilution protocols for feline tissues. A polymer detection system was applied as a secondary antibody for one hour, and immunoreactive cells were visualized via colorimetric detection (3,30-diaminobenzidine). A normal lymph node was used as a positive control, and isotype immunoglobulin at the same concentration of primary antibody was used as a negative control.

For immunohistochemistry analysis, samples were classified semiquantitatively for BAX, caspase-3 and VEGF-A and quantitatively for Ki67. For BAX, VEGF-A, and Caspase-3, the samples were categorized on the basis of intensity as follows: score 0 (no expression), score 1 (< 10% positive cells), score 2 (10% up to 50%), and score 3 (> 50%), according to the previous literature [19]. Samples with scores of 0 and 1 were considered to have low expression, and samples with scores of 2 and 3 were considered to have high expression. For Ki-67, three microphotographs at 40x magnification were taken, and the positively reacting cells were counted, with the average number of positive cells calculated across the three areas. The samples whose expression was not 50% or lower were considered to have low expression, and the samples whose expression was greater than 50% were considered to have high expression. This segregation was performed on the basis of previous literature [20].

### Analysis of Results and Statistics

The data were tabulated in an Excel (Microsoft 365, 2023) spreadsheet and analyzed for frequency and distribution. A normality test indicated a nonparametric distribution of the data. Survival analysis was conducted via Kaplan-Meier curves. For the Kaplan-Meier curve, the median expression of the markers was calculated, and the animals were subdivided into high- and low-expression groups on the basis of the description above. Fisher’s exact test was used to verify the associations between the semiquantitative immunohistochemical score and the clinical and pathological variables in a correlation matrix. Statistical analysis was performed via GraphPad Prism 8.0, with a p value considered significant at less than 0.05. For the statistical analysis, we separated the cats that received chemotherapy from those that did not receive any treatment.

## RESULTS

### Clinical information

Thirty cats met the inclusion criteria. In 24 of the 30 cases, the cats were mixed breed, and the average age of the patients was 10.6 years (±10). Among the 30 selected cases, we found 19 cats with information regarding infection with Feline Leukemia Virus (FeLV), of which 16 were negative for FeLV and three were positive. The remaining patients did not have information on FeLV status. The complete clinical information can be found in Supplementary Table 1.

**Table 1.** Comparative analysis of antibody expression, chemotherapy, and survival This table summarizes the relationships between antibody expression levels (low/high), chemotherapy treatment, and survival outcomes. Statistical significance is indicated by p values. The “Variable” in the survival (days) column denotes the range of observed survival times.

| Antibody | Expression Level | Chemotherapy | Survival (Days) | p value | Statistical Observation |
| --- | --- | --- | --- | --- | --- |
| BAX | Low | No | 0 - 500 | 32 | Significantly lower survival compared to high expression |
|  | High | No | 50 - 1500 |  |  |
|  | Low | Yes | 0 - 1000 | <0.0001 | Survival still lower in untreated compared to treated |
|  | High | Yes | up to 2000 |  |  |
| CASPASE 3 | Low | No |  | 222 | Trend toward lower survival in low expression |
|  | High | No |  |  |  |
|  | Low | Yes | 0 - 500 | <0.0001 | Survival lower in untreated compared to treated |
|  | High | Yes | up to 1750 |  |  |
| KI67 | Low |  |  | <0.0001 | Significantly lower survival in high expression |
|  | High |  |  |  |  |
|  | Low | Yes | up to 2000 | 184 | Survival lower in untreated compared to treated |
|  | High | Yes | 0 - 1000 |  |  |
| VEGF-A | Low |  |  | 556 | Not significant, trend to lower survival in high expr. |
|  | High |  |  |  |  |
|  | Low | Yes | up to 2000 | <0.0001 | Survival lower in untreated compared to treated |
|  | High | Yes | 0 - 1000 |  |  |

Treatment data were available for 24 cats, of which 17 underwent chemotherapy, and seven did not receive any chemotherapy. The first protocol included COP (cyclophosphamide, vincristine, and prednisone) for four patients, CHOP (doxorubicin, cyclophosphamide, vincristine, and prednisone) for three patients, chlorambucil combined with prednisone for nine patients, or corticosteroids alone for one patient. The mean overall survival of these patients was 137.5 (± 90) days.

### Histopathological and immunohistochemical analysis

With respect to histopathological classification, 17 patients were classified as lymphocytic lymphomas, and 13 were classified as lymphoblastic lymphomas. Immunophenotyping revealed that most lymphomas were of T-cell origin (22/30), whereas B-cell lymphomas accounted for 8/30, and in three cases, the cell origin could not be determined. On the basis of nuclear size and immunophenotyping, for 4/30 B lymphoblastic lymphomas, nine (9/30) T lymphoblastic lymphomas, 4/30 B lymphocytic lymphomas, and 13 (13/30) T lymphocytic lymphomas were identified.

Regarding immunohistochemistry, 73.3% (22/30) of the patients had high levels of BAX expression, and the remaining 26.7% (8/30) had low levels of BAX expression (Fig 1 A and B). For caspase-3, 70% (21/30) presented high expression, and 30% (9/30) presented low expression (Fig 1 C and D). Fifty percent (15/30) of the cases presented high Ki67 expression, and the remaining 50% (15/30) presented low expression (Fig 1 E and F). For VEGF-A, 50% (15/30) of the cases presented no low expression, and 50% (15/30) presented high expression (Fig 1 G and H).

**Fig 1.**
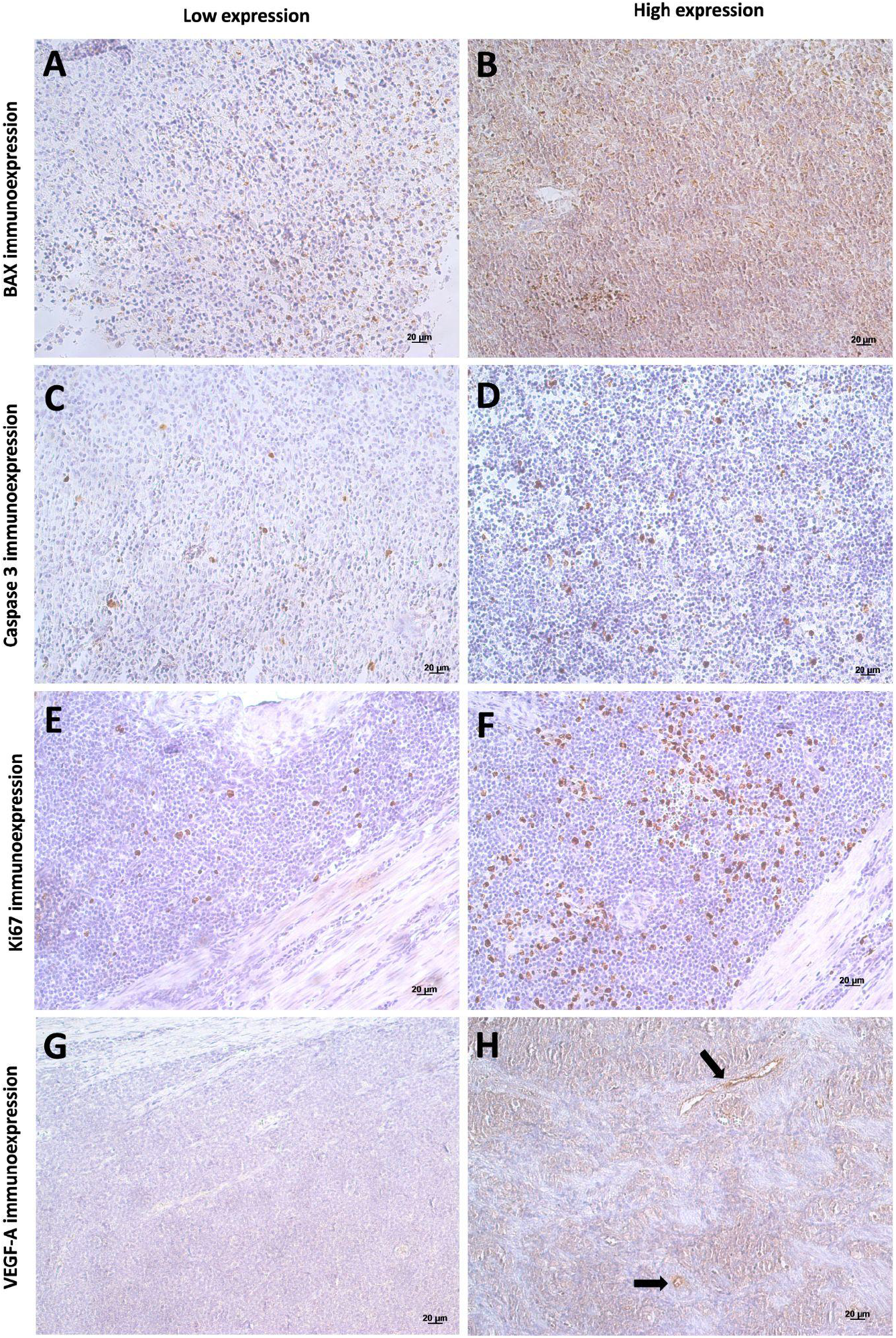
Low and high expression of BAX, Caspase-3, KI67 and VEGF. Immunoexpression of BAX, caspase-3, Ki-67, and VEGF-A in tumor samples with low (left column) and high expression (right column).

### Associations with clinical parameters

First, we evaluated the prognostic value of BAX, Caspase-3, VEGF-A, and Ki67 in feline intestinal lymphoma patients, independent of whether they received chemotherapy. Considering the variable histological subtypes (lymphocytic versus lymphoblastic), BAX immunoexpression (low versus high) did not differ (p > 0.9999). However, when we associated BAX immunoexpression with survival, we found a significant association. Cats with low levels of BAX expression were associated with shorter survival times (p = 0.0032) (Fig 2A). With respect to Caspase-3 expression, no significant difference was found when we compared the lymphoblastic and lymphocytic subtypes (p = 0.1776). On the other hand, Caspase-3 expression was significantly associated with overall survival (p = 0.0222) (Fig 2C). Patients with low Caspase-3 expression have a longer survival time.

**Fig 2.**
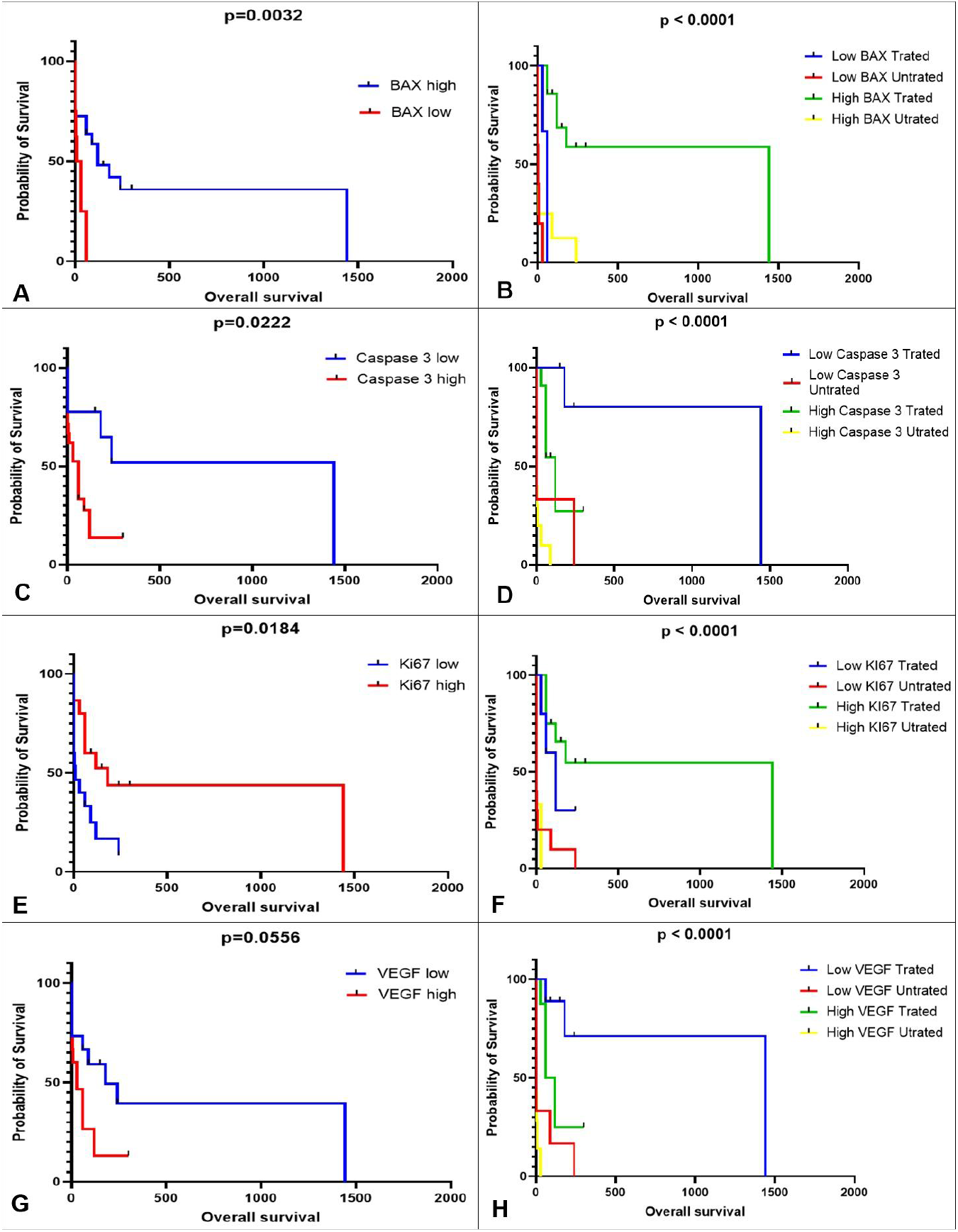
K-M survival curves according to the expression levels of BAX, caspase-3, Ki-67, and VEGF-A. (A, C, E, G) OS stratified by low and high expression of BAX, caspase-3, Ki-67, and VEGF-A, respectively. (B, D, F, H) OS according to expression level combined with treatment status.

VEGF-A scores also showed high variability between these lymphoma subtypes, and no significant difference was observed in VEGF-A immunoexpression between the lymphoblastic and lymphocytic subtypes (p = 0.3561). VEGF-A immunoexpression by neoplastic lymphocytes was not associated with overall survival (p = 0.0556) (Fig 2E). In terms of Ki67 expression, no significant difference was detected between cats with lymphoblastic lymphomas and those with lymphocytic lymphomas (p = 0.2309). However, we found an association between Ki67 expression and overall survival (p = 0.0184). Patients with high Ki67 expression experienced a shorter survival time (Fig. 2G).

Afterward, we separated our sample group according to the treatment received by the patient. When we combined this approach, the prognostic status of the biomarker assessed increased. First, we compared the overall survival of patients who received chemotherapy (independent of the protocol) with that of patients who did not receive any treatment. Chemotherapy significantly improved the overall survival of patients (p = 0.0008). When we compared biomarker expression segregated among patients who received or did not receive chemotherapy (i.e., low-BAX-treated patients, high-BAX-treated patients, low-BAX-treated patients, low-BAX-treated patients, and low-BAX-treated patients), we detected significant differences in all the markers. The individual results for all comparisons are presented in Table 1.

## DISCUSSION

The 30 selected animals originated from different locations, spanning the period from 2008--2023. Mixed-breed cats (MBCs) are predominant in most studies involving feline populations, and the results of this study are not different, with 80% of the subjects belonging to this category. A similar trend was observed in the Brazilian study by Cristo et al. [21], where mixed-breed cats accounted for 45 out of 53 evaluated cats [21]. The sex of the animals did not significantly affect the occurrence of lymphoma, as there were 11 males and 15 females. In a larger sample, these values tend to differ, as shown in the study by Sato et al., which included 163 cats, of which 92 were males and 71 were females. Nonetheless, the sex variable did not exhibit significance [22].

The literature indicates that cats have two age ranges with greater lymphoma recurrence: early adulthood and early old age [23]. Considering that felines become adults at two years of age and elderly at ten years of age, the cases presented in this study include patients within this proposed age range, with a median age of approximately ten years. The same median was reported in the study by Bik et al. [24].

With respect to the treatment of felines with lymphoma, a study conducted in the Netherlands in 2023 selected 174 felines with lymphoma, 110 of which were treated with the COP protocol [25]. A study by Collete et al. [26] examined the CHOP protocol in their patients. The patients in this study who received treatment utilized COP and CHOP, in addition to combinations of chemotherapeutics and corticosteroids. The felines included in this study had a mean overall survival of four months, regardless of whether they underwent treatment. Fabrizio et al. [27] reported a slightly longer survival time of eight months, whereas Limmer et al. [28] noted a shorter survival time of two months.

Feline leukemia virus (FeLV) was not investigated in all the cases in this study; however, 19 animals were tested for the virus, three of which had positive results, and all their lymphomas were of the alimentary type. In contrast, Limmer et al. presented findings in which six cats (6/26) tested positive for FeLV, with multicentric lymphoma being the dominant classification (2/6) [28].

Immunohistochemical labeling of Ki-67 is typically observed in the cell nucleus [29]. Rebollada-Merino et al. [30] reported in their study on intestinal lymphomas in felines that Ki-67 is expressed diffusely in the epithelial crypts. The same pattern was observed in the standardized samples used in this study. BAX belongs to the BCL family; intestinal lymphomas may exhibit intense expression of this biomarker, corroborating findings from our experiment [30] [31]. Caspase-3 exhibited sparse labeling, which can also be interpreted as granular, yielding results similar to those reported in the study by Silva et al. [30] on mammary carcinomas in felines. Finally, VEGF-A demonstrates the formation of new vessels in stromal tissues within neoplastic cells, a finding that was also reported by Chen et al. in their 2020 study [18].

When the labeling intensity of each biomarker used was analyzed, a relatively high concentration of samples that exhibited low or intermediate labeling was detected. For Ki-67, this finding indicates that the degree of cellular proliferation in these lymphomas was moderate or low. High levels of BAX may indicate a greater tendency toward apoptosis, whereas low levels of labeling could suggest a reduced ability to induce apoptosis, which may be related to increased cellular resistance, indicating the persistence of neoplastic cells [32]. Caspase-3 also measures apoptosis; however, low levels of labeling suggest potential resistance to apoptosis, which may be related to more aggressive lymphoma behavior [33]. Finally, low labeling of VEGF-A may indicate reduced induction of angiogenesis, leading to decreased tumor vascularization [34].

The high index of BAX in patients with intensity scores of 2 and 3 proved to be a good prognostic marker for feline lymphomas. A study published in BMC Cancer analyzed the expression of Bax as an independent prognostic marker in oral squamous cell carcinoma (OSCC). The results indicated that high expression of Bax was associated with a significant improvement in 5-year specific survival (85.7% for tumors with high expression versus 50.3% for those with low expression) [35]. In contrast, high expression of Caspase-3, VEGF-A, and Ki-67 has been associated with poorer survival in various lymphomas, highlighting their potential as markers of disease progression. In feline lymphomas, increased apoptotic activity, indicated by increased BAX expression, was linked to prolonged survival, whereas VEGF-A and Ki-67 did not directly correlate with prognosis [36]. In human studies, the expression of Caspase-3 and Ki-67 has been explored in ampullary carcinomas, where a negative correlation between Caspase-3 and Ki-67 was observed, suggesting their complementary roles in apoptosis and proliferation [37]. Furthermore, VEGF-A has been implicated in promoting angiogenesis and disease progression in various cancers, including lymphomas, although its prognostic value remains debated [38]. These findings underscore the need for a nuanced interpretation of these markers, considering the tumor type and biological context.

## CONCLUSION

It was concluded that BAX, Caspase-3 and Ki67 are potential prognostic markers in feline intestinal lymphomas, and future prospective studies are necessary for validation.

## Acknowledgments

This study was supported by the Coordenação de Aperfeiçoamento de Pessoal de Nível Superior – Brazil (CAPES) – Finance Code 001.

## Supporting Information

**Table S1.** Feline lymphoma cases. Age was calculated in years. For sex, “M” was used for males, and “F” was used for females. The breed was classified as “mixed” for felines without a defined breed and “Siamese” for Siamese cats. The size of the lymphocyte nucleus was classified as lymphocytic (smaller than a red blood cell) or lymphoblastic (larger than a red blood cell). Immunophenotyping results were divided into T-cell lymphoma and B-cell lymphoma. Chemotherapy treatment was separated into those who did not undergo chemotherapy (No) and those who did (Yes - 1. COP, 2. CHOP, 3. Chlorambucil and corticosteroids, 4. Corticosteroids). Survival was measured in days. The symbol (-) was used when no information was available.

| Cases | Age | Sex | Breed | Nucleus Size | Immunophenotype | Chemotherapy | Survival |
| --- | --- | --- | --- | --- | --- | --- | --- |
| Case 1 | 21 | M | Mixed | Lymphocytic | T | No | 90 |
| Case 2 | 9 | F | Siamese | Lymphoblastic | B | No | 0 |
| Case 3 | 11 | M | Mixed | Lymphocytic | T | Yes (4) | 30 |
| Case 4 | 13 | F | Mixed | Lymphocytic | T | No | 240 |
| Case 5 | - | - | Mixed | Lymphoblastic | B | - | 0 |
| Case 6 | - | F | Mixed | Lymphoblastic | T | - | 0 |
| Case 7 | 10 | M | Mixed | Lymphoblastic | T | No | 30 |
| Case 8 | 15 | F | Mixed | Lymphocytic | T | No | 3 |
| Case 9 | 12 | M | Mixed | Lymphocytic | B | - | 0 |
| Case 10 | - | - | Mixed | Lymphocytic | - | - | 0 |
| Case 11 | - | - | Mixed | Lymphocytic | B | - | 0 |
| Case 12 | - | - | Mixed | Lymphoblastic | T | - | 0 |
| Case 13 | 11 | F | Mixed | Lymphocytic | T | No | 4 |
| Case 14 | 10 | F | Mixed | Lymphoblastic | B | Yes (1) | 60 |
| Case 15 | 8 | M | Mixed | Lymphocytic | T | Yes (2) | 180 |
| Case 16 | 12 | M | Mixed | Lymphoblastic | T | Yes (1) | 60 |
| Case 17 | 7 | M | Mixed | Lymphoblastic | T | Yes (3) | 120 |
| Case 18 | 6 | F | Mixed | Lymphoblastic | B | Yes (2) | 60 |
| Case 19 | 1 | F | Mixed | Lymphoblastic | T | Yes (1) | 60 |
| Case 20 | 10 | F | Mixed | Lymphoblastic | T | Yes (3) | 240 |
| Case 21 | - | F | Mixed | Lymphoblastic | T | Yes (3) | 300 |
| Case 22 | 7 | M | Mixed | Lymphocytic | T | Yes (3) | 150 |
| Case 23 | 7 | M | Mixed | Lymphocytic | T | Yes (2) | 120 |
| Case 24 | 11 | F | Mixed | Lymphocytic | T | Yes (3) | 240 |
| Case 25 | 15 | M | Mixed | Lymphoblastic | T | Yes (3) | 90 |
| Case 26 | 15 | F | Mixed | Lymphocytic | T | Yes (3) | 1440 |
| Case 27 | 12 | F | Mixed | Lymphocytic | B | Yes (3) | 240 |
| Case 28 | 11 | M | Mixed | Lymphocytic | T | No | 10 |
| Case 29 | 10 | F | Mixed | Lymphocytic | B | Yes (1) | 60 |
| Case 30 | 11 | F | Mixed | Lymphocytic | T | Yes (3) | 300 |

## REFERENCES

1. Gabor LJ, Canfield PJ, Malik R. Clinical and anatomical features of lymphosarcoma in 118 cats. Aust Vet J. 2016;78(6):412–417.

2. Marconato L, Gelain ME, Comazzi S. The dog as a possible animal model for human non-Hodgkin lymphoma: A review. Hematol Oncol. 2019;27(2):53–64.

3. Noronha LF, Cristo TG, Biezeus G, Furlan LV, Costa LS, Pereira Lhhs, et al. Caracterização Anatopatológica dos Linfomas em Gatos Domésticos. In: 28° SIC UDESC; 2018.

4. Nogueira MM, Melo MM. Linfoma alimentar linfocítico felino – Uma revisão de literatura. Rev Bras Hig Sanid Anim. 2020;14(3):1–15.

5. Silva DHL, Ecco R, Pierizan F, Cassali GD, Reis JKP, Gonçalves ABB, et al. Classification of lymphoma in cats and its relationship with detection of feline leukemia virus proviral DNA. Pesq Vet Bras. 2022;42:07021.

6. Duffy MJ, Crown J. Biomarkers for Predicting Response to Immunotherapy with Immune Checkpoint Inhibitors in Cancer Patients. Clin Chem. 2019;65(10):1228–1238.

7. Smith RA, Tudor-Smith C, Neoptolemos JP, Ghaneh P. Meta-analysis of immunohistochemical prognostic markers in resected pancreatic cancer. Br J Cancer. 2011;104:1440–1451.

8. Gouveia GPF. Avaliação da Expressão Imunohistoquímica de EGFR e Ki-67 no Carcinoma de Células Escamosas Oral em Gato [dissertação]. Lisboa: Faculdade de Medicina Veterinária da Universidade de Lisboa; 2022.

9. Gadbail AR, Sarode SC, Chaudhary MS, Gondivkar SM, Takade SA, Yuwanati M, et al. Ki67 Labeling Index predicts clinical outcome and survival in oral squamous cell carcinoma. J Appl Oral Sci. 2021;29:e20200751.

10. Youle RJ, Strasser A. The BCL-2 protein family: opposing activities that mediate cell death. Nat Rev Mol Cell Biol. 2008;9(1):47–59.

11. Smith A, Wiseman DH, Liao Z, Bell AK, Hasserjian RP, Wood BL. Expression of Bax in Diffuse Large B-Cell Lymphoma: Prognostic Significance and Relationship with Tumor Microvessel Density and the Apoptosis Index. Clin Cancer Res. 2009;15(4):1278–87.

12. Ho LH, Read SH, Dorstyh L, Lambrusco L, Kumar S. Caspase-2 is required for cell death induced by cytoskeletal disruption. Oncogene. 2008;27(51):7114–22.

13. Shibuya M. Vascular endothelial growth factor (VEGF) and its receptor (VEGFR) signaling in angiogenesis: a crucial target for anti- and pro-angiogenic therapies. Genes Cancer. 2011;2(12):1097–1105.

14. Pohlman LM, Higginbotham ML, Welles EG, Johnson CM. Immunophenotypic and histologic classification of 50 cases of feline gastrointestinal lymphoma. Vet Pathol. 2009;46(2):259–68.

15. Madewell BR, Gandour-Edwards R, Edwards BF, Matthews KR, Griffey SM. Bax/bcl-2: cellular modulator of apoptosis in feline skin and basal cell tumors. J Comp Pathol. 2001;124(2-3):115–21.

16. Macente BI, Apparicio M, Mansano CFM, Tavares MR, Fonseca-Alves CE, Sousa BP, et al. Effect of cryopreservation on sperm DNA fragmentation and apoptosis rates in the testicular tissue of domestic cats. Anim Reprod Sci. 2019;211:106224.

17. Soares M, Ribeiro R, Carvalho S, Peleteiro M, Correia J, Ferreira F. Ki-67 as a Prognostic Factor in Feline Mammary Carcinoma: What Is the Optimal Cutoff Value? Vet Pathol. 2016;53(1):37–43.

18. Chen B, Lin SJ, Li WT, Chang HW, Pang VF, Chu PY, Lee CC, et al. Expression of HIF-1α and VEGF in feline mammary gland carcinomas: association with pathological characteristics and clinical outcomes. BMC Vet Res. 2020;16(1):125.

19. Peiró G, Diebold J, Baretton GB, Kimmig R, Löhrs U. Cellular apoptosis susceptibility gene expression in endometrial carcinoma: correlation with Bcl-2, Bax, and caspase-3 expression and outcome. Int J Gynecol Pathol. 2001;20(4):359–67.

20. Freiche V, Paulin MV, Cordonnier N, Huet H, Turba ME, Macintyre E, et al. Histopathologic, phenotypic, and molecular criteria to discriminate low-grade intestinal T-cell lymphoma in cats from lymphoplasmacytic enteritis. J Vet Intern Med. 2021;35(6):2673–2684.

21. Cristo TG, Biezeus G, Noronha LF, Pereira Lhhs, Withoeft JA, Furlan LV, et al. Feline Lymphoma and High Correlation with Feline Leukemia Virus Infection in Brazil. J Comp Pathol. 2019;166:20–28.

22. Sato H, Yasuhito F, Goto-Koshino Y, Uchida K, Ohno K, Tsujimoto H. Prognostic Analyses on Anatomical and Morphological Classification of Feline Lymphoma. J Vet Med Sci. 2014;76(6):807–811.

23. Nelson RW, Couto CG. Medicina interna de pequenos animais. Rio de Janeiro: Elsevier Editora Ltda; 2015.

24. Bike CA, Ruijter BR, van den Bossche L, Teske E, Niessen SJM, Forcada Y. Pegylated-L-asparaginase therapy for feline large cell lymphoma: 82 cases (2017-2020). J Feline Med Surg. 2023;1–9.

25. Versteegh H, Zandvliet MMJM, Feenstra LR, van der Steen FEMM, Teske E. Feline Lymphoma Patient Characteristics and Response Outcome of COP-Protocol in Cats with Malignant Lymphoma in The Netherlands. Animals. 2023;13(16):2667.

26. Collete SA, Allstadt EM, Chon W, Vernau AN, Smith LD, Garrett K, et al. Treatment of feline intermediate-to high-grade lymphoma with a modified university of Wisconsin–Madison protocol: 119 cases (2004–2012). Vet Comp Oncol. 2015;14(S1):136–146.

27. Fabrizio F, Calam AE, Dobson JM, Middleton SA, Murphy S, Taylor SS, et al. Feline mediastinal lymphoma: a retrospective study of signalment, retroviral status, response to chemotherapy and prognostic indicators. J Feline Med Surg. 2014;16(9):637–644.

28. Limmer S, Eberle N, Nerschbach V, Nolte I, Betz D. Treatment of feline lymphoma using 12-week, maintenance-free combination chemotherapy protocol in 26 cats. Vet Comp Oncol. 2014;14(S1):21–31.

29. Silva MN, Leite JS, Mello MFV, Silva Kvgc, Corgozinho KB, Souza HJM, et al. Histologic evaluation of Ki-67 and cleaved caspase-3 expression in feline mammary carcinoma. J Feline Med Surg. 2017;19(4):440–445.

30. Rebolla-Merino A, Porras N, Calvo-Ibbitson A, Rodrigues-Franco F, Rodrigues-Bertos A. Bcl-2 Immunoexpression in Feline Epitheliotropic Intestinal T-Cell Lymphomas. Vet Sci. 2022;9:168.

31. Youle RJ, Strasser A. The BCL-2 protein family: opposing activities that mediate cell death. Nat Rev Mol Cell Biol. 2008;9(1):47–59.

32. Moany M, Ahmed MM, Al-Rejaie SS. The Role of NF-κB and Bax/Bcl-2/Caspase-3 Signaling Pathways in the Protective Effects of Sacubitril/Valsartan (Entresto) against HFD/STZ-Induced Diabetic Kidney Disease. Biomedicines. 2022;10:2863.

33. Loureiro LVM, Neder L, Callegaro-Filho D, et al. The immunohistochemical landscape of the VEGF family and its receptors in glioblastomas. Surg Exp Pathol. 2020;3:9.

34. Suter S. Feline gastrointestinal lymphomas: diagnosis and treatment. In: Veterinary Meeting & Expo; 2023; Orlando, FL. Proceedings. Orlando: DVM360; 2023.

35. Narayan K, et al. BAX expression measured by AQUAnalysis is an independent prognostic marker in oral squamous cell carcinoma. BMC Cancer. 2020;20(1):1–12.

36. Silva RT, Bombonato JH, Rocha NS, et al. Prognostic evaluation of Ki-67, VEGF, and BAX expression in feline lymphomas: Insights into cellular proliferation, apoptosis, and angiogenesis. UNESP Repository. 2024.

37. Ozluk Y, Kilicaslan I, Durak H, et al. Expression of livin, Caspase-3, and Ki-67 in ampullary carcinomas and their correlation with clinicopathological parameters. PubMed Central. 2013.

38. De Nardi AB, Rodriguez MI, Brunetto MA, et al. COX-2, VEGF, and Caspase-3 immunoreactivity in canine mammary tumors: Correlation with biological behavior. Brazilian Digital Library of Theses and Dissertations. 2016

